# Differences in player position running velocity at lactate thresholds among male professional German soccer players

**DOI:** 10.1101/592188

**Authors:** René Schwesig, Stephan Schulze, Lars Reinhardt, Kevin G. Laudner, Karl-Stefan Delank, Souhail Hermassi

## Abstract

This study investigated the differences in running velocities at specific lactate thresholds among male German soccer players. One hundred fifty-two professional (3^rd^ league: n=82; 4^th^ league: n=70) male soccer players (mean ± SD; age: 24.7 ± 4.38 years, body mass: 80.7 ± 7.36 kg, body height: 1.83 ± 0.06 m) volunteered for the investigation. Players were categorized as goalkeepers, central defenders, central midfielders, wings and forwards. Players completed a treadmill test, at incremental speeds, to determine running velocity at different blood lactate concentrations (v2=2 mmol/l; v4=4 mmol/l; v6=6 mmol/l). Results indicate that, wings displayed the lowest body mass (76.2 ± 6.08 kg) and body height (1.79 ± 0.06 m). In contrast, goalkeepers were the tallest athletes in the whole sample (1.90 ± 0.03 m), forwards were the heaviest players (85.4 ± 6.03 kg). In addition, we detected the largest difference between positions for running velocity at the lactate threshold v2 (p=0.002). The running data revealed that only the goalkeepers had significantly lower velocities at the lactate thresholds compared to the field players. The central midfielders showed the highest performance level at the lactate thresholds (v2: 12.5 ± 1.20 km/h; v4: 15.2 ± 1.14 km/h; v6: 16.6 ± 1.13 km/h). In conclusion, this study provides soccer and position-specific reference data for the performance of male professional German soccer players in order to evaluate the running performance in a valid way. In this context, it is necessary to extend the database for the second and first league. Furthermore, it is important to assess the running performance during competition matches over the entire season in order to validate the endurance test performance data.

## Introduction

Soccer is one of the most widely played and complex sports in the world, where players need technical, tactical, and physical skills to succeed. However, studies to improve soccer performance have often focused on technique and tactics at the expense of physical resources such as endurance, strength, and speed [1–3].

Endurance in soccer is characterized as low to high-intensity intermittent running performance [4,5] and is represented by the physical amount of work carried out throughout a match [6]. Furthermore, it has been observed that soccer players reach peak running speeds close to 32 km⋅h^−1^ during match-play [7,8]. This quality of sprinting speed depends on several factors including the level of practice and the players’ age [9–10]. Indeed, it has been shown that elite players are faster during the first 10 m of a 30 m sprint test than amateurs [10] and that older players are faster covering a 40 m sprint test in highly trained young soccer players [11].

A number of field and laboratory tests are currently used in elite soccer in order to evaluate training status of the players, to predict match performance, and to determine the effect of training [12]. Various incremental exercise tests and intermittent shuttle run tests are currently used to evaluate training status and training adaptations, as well as to predict running performance during matches in soccer players [12, 13].

Incremental exercise tests are capable of providing input into aerobic endurance performance parameters [14–17]. In addition, heart rate and lactate thresholds are commonly used as a sensitive indicator of changes in training status in soccer players [18].

No previous work has investigated differences in running performance among different playing positions in male professional soccer players. This information is important in order to generate evidence based implications for the soccer specific endurance training. Based on the results of Foehrenbach et al. [19] a running speed of 4 m/s is often used at the 4 mmol/l lactate threshold to judge the running performance of soccer players [20–23]. We hypothesized that running velocity at various lactate thresholds are significantly different by playing position in male soccer players.

## Methods

### Participants

The test data were collected as part of performance diagnostic measures in the period between 2015 and 2018 and resulted from tests with teams from the 3^rd^ and 4^th^ German soccer leagues. A total of 304 data sets were available for the evaluation (3^rd^ league: n=151; 4^th^ league; n=153). In order to avoid correlated observations, we calculated the median from different tests from the same player for every parameter. After this calculation, 152 professional (3^rd^ league: n=82; 4^th^ league: n=70) male soccer players (mean ± SD; age: 24.7 ± 4.38 years, body mass: 80.7 ± 7.36 kg, body height: 1.83 ± 0.06 m) were included in the statistical analysis. All players were healthy and completed the tests following a day of rest. Written informed consent was given after explanation of the study design and potential risks. In addition, study participants were informed that they could withdraw from the project at any time without penalty. To provide an in-depth analysis of team soccer, results were analyzed for the entire group and according to individual playing positions. This study was carried out in accordance with the recommendations of institutional human research review board at the Department of Orthopedic and Trauma Surgery, (UKH) Halle-Wittenberg, committee. The (UKH) Halle-Wittenberg, committee approved the protocol. All subjects gave written informed consent in accordance with the Declaration of Helsinki.

### Experimental Design

This study examined if differences in running velocity at specific lactate thresholds exist between male soccer players of the third league (n=151) and fourth league (n=153) and playing positions (Figure 1). Between leagues, we found no relevant differences. Therefore, we compared only between the positions.

**Figure 1:**
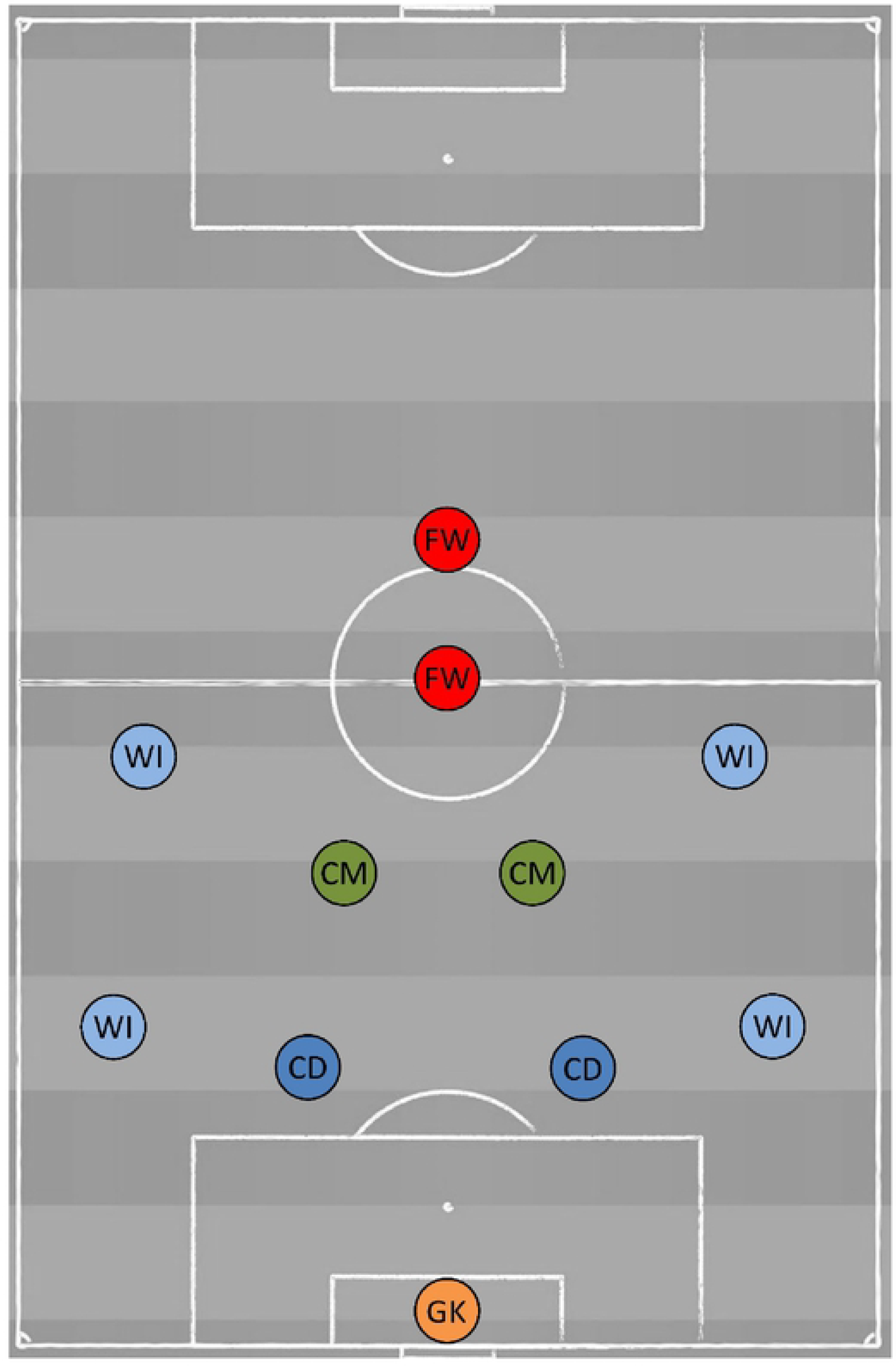
Soccer playing positions: goalkeepers (GK), central defenders (CD), central midfielders (CM), wings (WI), forwards (FW)

The subjects were carefully familiarized with the testing protocol, as they had been previously tested on several occasions in season for training prescription purposes. All of the players within a given team were assessed on the same day, and the tests were performed in the same order.

All tests were conducted at the start (May and June 2015 - 2018) of the pre-seasonal training. To reduce the influence of uncontrolled variables, all participants were instructed to maintain their typical lifestyle and diet habits before and during the study. Subjects were told not to exercise on the day ahead of a test and to consume their last (caffeine-free) meal at least 3 h before the scheduled testing. Additionally, they drank at least 0.5 liter of pure water during the last hour before testing. Regular sleep ahead of the protocol was also requested. During all performance-based testing, athletes were instructed to perform at their maximum ability. The local medical ethics committee of the medical faculty approved the study (reference number: 2013-13).

### Anthropometry measurement

Body height was measured using a stadiometer (Bodymeter 206, SECA®, Hamburg, 164 Germany to 0.1 cm) and body mass on an electrical scale (InBody120, model 165 BPM040S12FXX, Biospace, Inc., Seoul, Korea, to 0.1 kg).

### Resting measurements

Heart rate monitors (Polar Team Pro; Polar Electro Oy, Kempele, Finland) were fitted to the players 15 minutes before the start of the treadmill test. Heart rates (HR) were monitored using short-range telemetry with a one second recording interval. Athletes were asked to sit in a comfortable position for a duration of at least ten minutes without speaking. We measured heart rate before, during and after the treadmill test and determined the minimum heart rate using a ten seconds interval for calculation of the resting heart rate before the treadmill test. The measuring of heart rate was only necessary in order to monitor the test and calculate heart rate ranges for endurance training.

### Treadmill test

Athletes completed a treadmill test at incremental speeds, either in laboratory or gym, to determine running velocity at specific lactate threshold levels. The test started at a speed level of 7.2 km/h, and an increment of 1.8 km/h every 3 min. The slope was set at 0%. Prior to testing, all subjects completed a two minutes treadmill running warm up at a speed of 7.2 km/h in order to adapt to running on the treadmill.

We measured the lactate concentration in hemolyzed whole blood using an enzymatic lactate analyzer (Super GL easy; Dr. Müller Gerätebau GmbH, Freital, Germany) to determine three running levels. These three levels consisted of: v2=2 mmol/l lactate threshold, v4=4 mmol/l lactate threshold, and v6=4 mmol/l lactate threshold. Blood samples (10 μl capillary blood) were collected from an athlete`s ear lobe. Afterwards test data was analysed (winlactat, mesics, Münster, Germany) in order to determine the running velocity at the v2, v4 and v6. Mathematical modelling of the lactate-velocity function was determined by either an exponential or a polynomial function, depending on the best fitting results (highest explained variance of the regression function).

### Statistical Analysis

Prior to descriptive and inference statistical analyses, the data was checked for multiple player entries. If there was more than one data set per player, the median of the test results was calculated and included into the analysis. Results were analysed for the entire group and for individual playing positions (wings, central defenders, central midfielders, forwards and goalkeepers). Descriptive statistics (mean, median, standard deviation, percentile 10, 25, 50, 75, 90) were ascertained for all parameters. Mean differences between positions were tested using a one-factorial (position) univariate general linear model. Differences between means were considered as statistically significant if p values were <0.05 and partial eta squared (η_p_^2^) values were higher than 0.10 [24].

All statistical analyses were performed using SPSS version 25.0 for Windows (SPSS Inc., Chicago, IL, USA).

## Results

### Anthropometric data

The investigated soccer players (24.7 ± 4.38 years; Table 1) displayed significant differences between positions for body height (p<0.001; η_p_^2^=0.375) and body mass (p<0.001; η_p_^2^=0.266) (Table 2). Wings showed the lowest body mass (76.2 ± 6.08 kg) and body height (1.79 ± 0.06 m) (Table 2), especially compared to goalkeepers and central defenders (body height and mass: p<0.001) and central midfielders (body mass: p=0.036).

In contrast, goalkeepers were the tallest athletes (1.90 ± 0.03 m), whereas forwards were the heaviest players (85.4 ± 6.03 kg). Age, body mass, resting heart rate and resting lactate were not significantly different between the five positions (Table 2).

### Performance data

The running data (Table 3, 4, 5) revealed that only the goalkeepers have a significantly lower endurance performance compared to the field players, especially to the central midfielders (v2: p=0.002 (Figure 2); v4: p=0.009 (Figure 3); v6: p=0.025 (Figure 4)).

**Figure 2:**
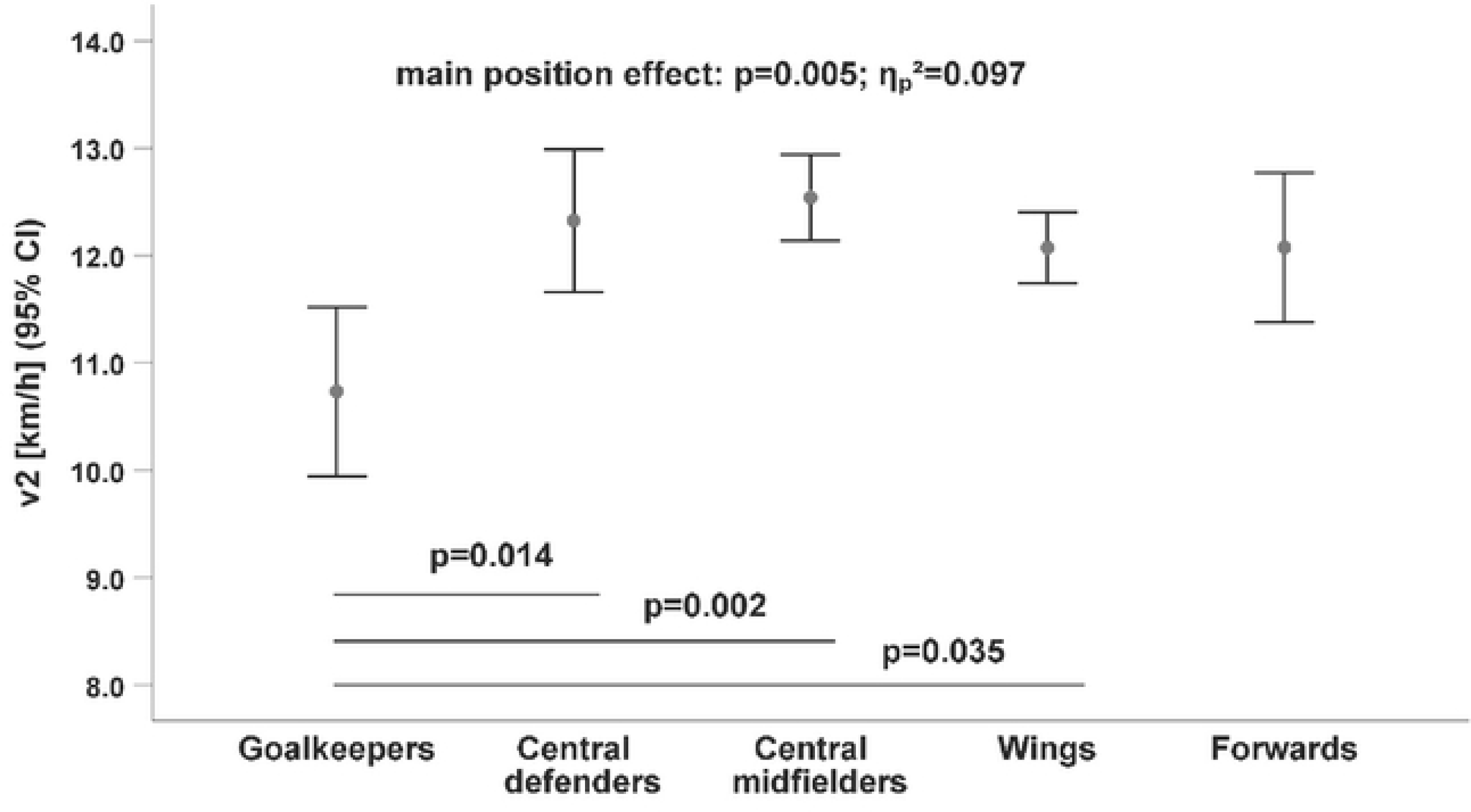
Differences in running velocity by playing position at the 2 mmol/1 lac tate threshold (v2). CI=confidence interval.

**Figure 3:**
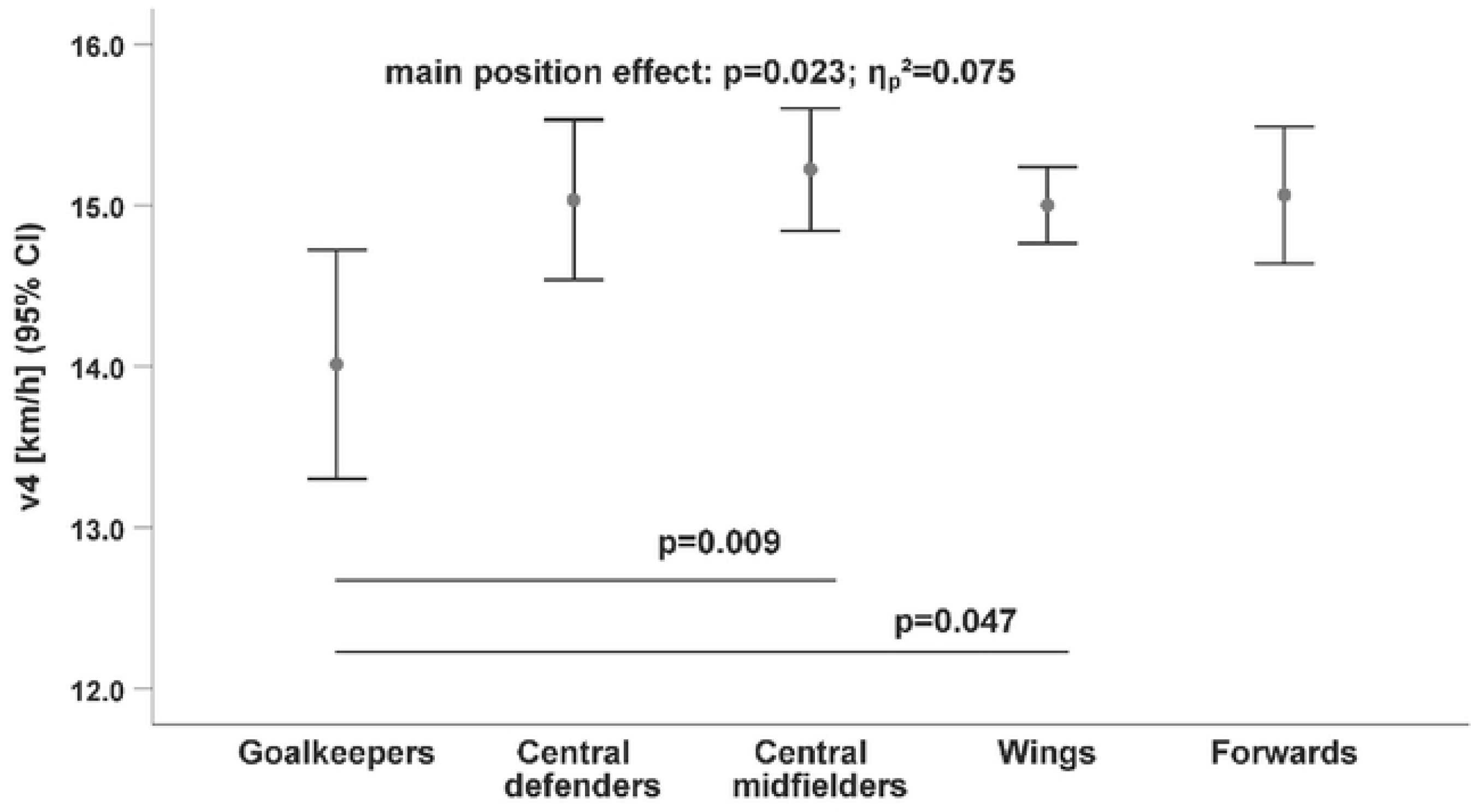
Differences in running velocity by playing position at the 4 mmol/1 lac tate threshold (v4). CI=confidence interval.

**Figure 4:**
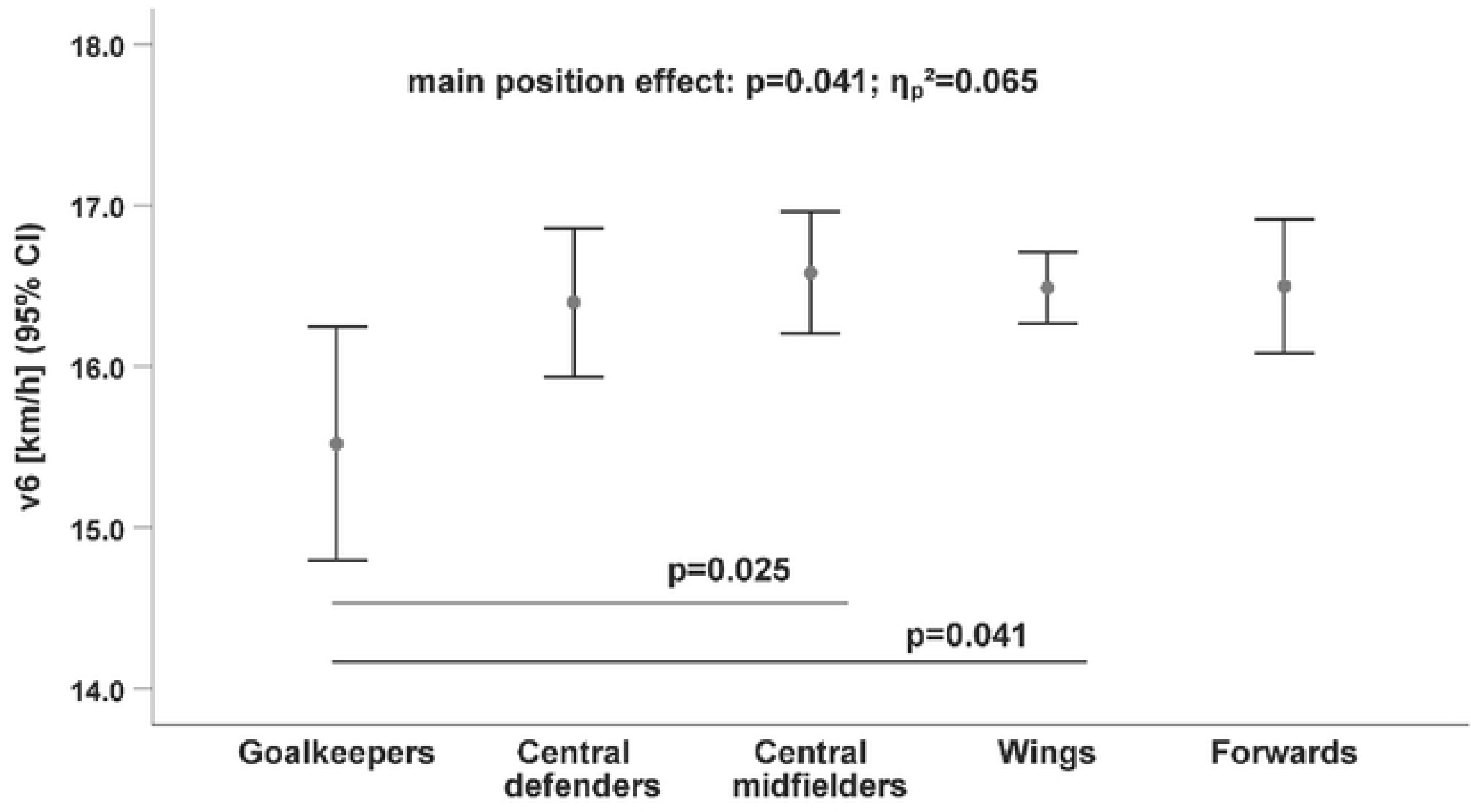
Differences in running velocity by playing position at the 6 mmol/1 lac tate threshold (v6). CI=confidence interval.

The difference between goalkeepers and central midfielders for the lactate threshold v2 (p=0.002; Figure 2) was the largest difference between positions. The endurance performance of the wings were also significantly higher at all lactate thresholds (v2: p=0.035; v4: p=0.047; v6: p=0.041; Table 6) compared to the goalkeepers.

There were no significant differences among the field players (Figures 2, 3, 4). Whereas, the goalkeepers were the players with the lowest performance level at all lactate thresholds (v2: 10.7 ± 1.17 km/h, p=0.005/η_p_^2^=0.097; v4: 14.0 ± 1.06 km/h, p=0.023/η_p_^2^=0.075; v6: 15.5 ± 1.08 km/h, p=0.041/η_p_^2^=0.065; Table 6), the central midfielders showed the highest performance level at the lactate thresholds (v2: 12.5 ± 1.20 km/h; v4: 15.2 ± 1.14 km/h; v6: 16.6 ± 1.13 km/h; Table 6).

## Discussion

The aim of this study was to assess playing position differences in running performance among male German soccer players. Our findings substantiate the hypothesis that different playing positions have specific running-related performance requirements among soccer players from the third and fourth German leagues.

In the present study, we detected several performance differences between playing positions, especially between the field soccer players and the goalkeepers. While the goalkeepers always had the lowest running velocity at all lactate thresholds, the central midfielders were the field players with the highest running performance. As expected, soccer players also showed significant differences in anthropometric characteristics, depending on their playing positions.

Although investigations of running performance have been previously conducted by other authors [18–23], comparability of these studies is difficult and limited due to differences in methodologies. Additionally, a classification deviating from our position assignment were provided by Bush et al. [25] and Suarez-Arrones et al. [26]. In contrast, to the present study, wing players were assigned to tactical role and tasks (wide defender, wide midfielder). Therefore, we chose the classification in our investigation, because the running requirements of the wing players (defensive and offensive) are very similar to each other [7] and therefore this classification seemed to be the most sensible and valid.

Rampinini et al. [7] investigated the match performance of professional soccer players using a computerized, semi-automatic video match analysis (speed, distance). Midfielders were the players with the highest running performance during the match. For example, midfielders covered 4, 13 and 15% more total distance than the fullbacks, forwards and centre-backs, respectively. The highest values recorded for time spent in the high-speed running zone (19.8-25.2 km/h) were performed by fullbacks and midfielders. These results correspond with our results, according to which the central midfielders were the players with the highest running performance.

The quality of lactate threshold data depends partly on the degree of exhaustion, standardization of the subsequent load [27], or on the kinetics of the test curve [28,29]. Contrary, evaluating fixed lactate thresholds, especially the 2 mmol/l lactate threshold, can be a disadvantage for fast player types. Some of these players produce high initial lactate values, but they also form higher maximum values at full load, in comparison with players who tend to have more slow twitching muscle fibers. Longitudinal investigations with soccer players from the 3^rd^ Greek league [22] showed a clearly lower performance at the 4 mmol/l lactate threshold (12.3-13.7 km/h) with a similar test design (3 min per speed level, with 2 km/h increment). Soccer players of the first three Greek leagues were examined with an almost identical test design to the one used in the current study, but lactate threshold was determined using the Dmax model [18]. Since threshold values in the Ziogas study ranged from 3.8 - 3.9 mmol/l, the comparison with the v4 from our study is possible. The players had an average threshold performance of 13.2 km/h (1^st^ league) to 12.3 km/h (3^rd^ league), which was also strongly different to our findings (below percentile 10: 13.7 km/h).

Another investigation examined the threshold performance of British youth soccer players [23]. The test design of this study differed significantly from the design of our investigation (4 min per speed level, with 0.5 km/h increment). During the season v4 speed of the players ranged between 13.6±0.3 km/h and 14.7±0.2 km/h. However, with the exception of preseason performance (13.6 km/h), the other reported values (14.7 km/h) were on a similar level to our data (P50: 15.0 km/h).

Altmann et al. [30] analyzed the physical capacity of 28 male soccer players (second German league). An incremental treadmill test with a similar test design (start speed: 6 km/h, 3 min per speed level, with 2 km/h increment) were used within a performance testing battery. One of the key parameters for endurance performance was the v4 lactate threshold. The authors reported almost identical results at v4 (average speed: 15.1 km/h, position-independent) for the field players compared with our sample (P50: 15.0 km/h). Obviously, the difference in league (third vs. second) is not a distinguishing criterion.

### Limitations

As in any scientific work, some methodological aspects need to be discussed. Due to the support of different teams in different cities, the implementation took place in local fitness studios with the treadmills available there. The test equipment was serviced in the usual maintenance intervals and met the technical specifications specified by the manufacturer. However, some differences in treadmill efficiency could have been present between tests.

To the best of our knowledge, there are no uniform guidelines for the running diagnostics of soccer players. Depending on the author collective, the tests for evaluating endurance performance are carried out as field tests on the track or alternatively on the treadmill. The load time per speed level varies between three and five minutes (treadmill) or up to 2.400 m per load distance (400 m track) [19, 21, 22]. Therefore, the discussion of the different results is difficult and conclusions should be drawn with great caution.

In German-speaking countries, a value of 4 m/s at 4 mmol/l lactate threshold is considered the gold standard for evaluating running performance of soccer players. From a methodological point of view, it should be noted that this value was initially calculated on the basis of running loads of 2.323 m per load level using a sample size of only thirty players [19]. All newer studies known to us work with significantly shorter step distances, so that the classification based on this value is likely to overestimate the performance of soccer players.

## Conclusions

Our results offer valuable information for soccer players, coaches, and technical staff by providing a valid classification of running performance by playing position. Soccer coaches may also benefit from these results in the preparation of individualized training derivations. Future research is needed to clarify the validity of these data and to compare test data with actual soccer match data.

## Acknowledgments

Special thanks to all soccer players and coaches for their valuable help and involvement in this study.

